# Bacterial Diversity in Feces of Wild Bald Eagles, Turkey Vultures and Common Ravens from the Pacific Northwest Coast, U.S.A

**DOI:** 10.1101/511147

**Authors:** Rocio Crespo, Scot E Dowd, Daniel E. Varland, Scott Ford, Thomas E. Hamer

## Abstract

Birds harbor diverse microorganisms in their guts, which collectively fulfill important roles in providing their hosts with nutrition and protection from pathogens. Although numerous studies have investigated the presence of certain pathogenic bacteria in the feces of wild birds, only a few have attempted to investigate the microbiota of the gut. This study analyzed the avian bacteria present in the cloaca of avian scavengers captured on coastal beaches of Washington and Oregon between 2013 and 2015: 10 turkey vultures (*Cathartes aura*), 9 bald eagles (*Haliaeetus leucocephalus*), and 2 common ravens (*Corvus corax*). We used illumina sequencing based on the V4 region of the 16s gene was to characterize the bacterial diversity. Our investigation revealed phylum-level differences in the microbiome of turkey vultures, compared with bald eagles and common ravens. Substantial microbiome differences were found between bald eagles and ravens below the phylum level. Although little is known about the possible relations among these microorganisms, our analyses provides the first integrated look at the composition of the avian microbiota and serves as a foundation for future studies in this area.

## Introduction

Birds harbor diverse microorganisms in their gut, which collectively fulfill important roles in providing their hosts with nutrition and protection from pathogens [1]. Gut microorganisms have been isolated and characterized by a variety of methods, such as culture-based assays, culture-independent DNA sequencing, and fluorescence *in situ* hybridization targeting the 16S rRNA gene [2]. Most of these studies are limited in scope and are focused on pathogenic bacteria such as *Salmonella*, *Escherichia coli*, *Campylobacter*, and *Clostridium perfringens* [3–5].

In recent years, metagenomics using next-generation sequencing has emerged as a way to analyze complex microbial communities and their functions [6]. This approach is capable of sequencing thousands or millions of amplified DNA molecules in a single run and, for bacteria, does not require conventional cloning and amplification. Additionally, metagenomic studies offer a powerful tool for the comprehensive and unbiased assessment of microbial diversity within the complex gut ecosystem by allowing examination of organisms not easily cultured in the laboratory [6, 7].

Few studies have used genetic sequencing to explore the intestinal microbiota of wild birds [8]. Furthermore, these studies reported gene sequences to phylum or class levels only, with no attempt to understand extant microorganism populations at the species level. In this study, we analyze the avian microbiome present in the cloaca of bald eagles, turkey vultures, and common ravens collected in coastal Washington and Oregon, U.S.A. Because research has demonstrated that intestinal microorganisms can influence the lifecycle and reproduction of their hosts [9], data presented in this paper can be important for a better understanding of the health and nutrition of these apex scavengers that comprise key components of the food web of the Pacific Northwest coast.

## Materials and methods

We captured avian scavengers in coastal Washington and Oregon with a carcass-baited net launcher [10]. We captured turkey vultures and common ravens from May through August and bald eagles from February through June. Once in hand, we marked birds for identification with visual identification leg bands (eagles and ravens) or patagial tags (vultures) [11].

### Aging

We assigned captured birds to two age classes, adult and immature. We classified bald eagles as adults (≥ 4 years old) with attainment of fully white head and tail plumage, yellow beak and yellow iris [12]. We classified turkey vultures as adults (≥ 2 years old) with attainment of a bright ivory bill [13]. Common ravens were ranked as adults (≥ 1 years old) by having glossy black wing and tail feathers and as immatures by having dull black or brownish wing and tail feathers [14].

### Sexing

When possible, we determined the sex of individuals using measurements. We sexed bald eagles using bill depth and hallux length measurements [15] and common ravens using foot pad length and body mass [16]. Using this approach, the sex of one raven captured was equivocal. In this case, we classified the individual as female because we observed it with another raven that we assumed was male based on its demonstrably larger size[17]. We did not sex captured turkey vultures because it is not possible to differentiate gender using plumage characteristics or measurements [13].

### Fecal sampling

We extracted up to 2 ml of fecal material directly from the cloaca of each bird using a disposable 3 ml pipette. In the field, we transferred fecal samples to cryovials and then placed them in a pre-frozen Golden Hour Thermal Container (www.credothermal.com). With no more than 8 hours of elapsed time, samples were stored at −20°C for up to one month and then at −70°C until processed in 2017.

Genetic sequencing of all 21 cloaca flora samples was conducted by MR DNA, Molecular Research, LP (www.MrDNAlab.com; Shallowater, Texas). In summary, the 16S rRNA gene V4 variable region PCR primers 515/806 were used in a single-step 30 cycle PCR using the HotStarTaq Plus Master Mix Kit (Qiagen, USA) under the following conditions: 94°C for 3 minutes, followed by 30 cycles of 94°C for 30 seconds, 53°C for 40 seconds and 72°C for 1 minute, after which a final, 5-minute elongation step at 72°C was performed on a MiSeq system (Illumina Inc, San Diego, California). After amplification, PCR products were checked in 2% agarose gel to determine the success of amplification and the relative intensity of bands. Samples were pooled and purified using calibrated Ampure XP beads (Beckman Coulter, Life Sciences, Indianapolis, Indiana). This purified PCR product was then used to prepare an Illumina DNA library (https://www.illumina.com/techniques/sequencing/ngs-library-prep.html).

Sequence paired ends were processed using MR DNA taxonomic analysis pipeline (www.MrDNAlab.com). In summary, the process included depletion of barcodes and primers from genetic sequences, paired sequences were merged, followed by removal of sequences <150bp, and then removal of sequences with ambiguous base calls and homopolymer runs exceeding 6bp. Following these steps, operational taxonomic units (OTUs) were generated at 97% homology and chimeras were removed. Final OTUs were taxonomically classified using BLASTn (https://blast.ncbi.nlm.nih.gov/Blast.cgi) against a database derived from Ribosomal Database Project II (http://rdp.cme.msu.edu) and the National Center for Biotechnical Information (www.ncbi.nlm.nih.gov). When a given sequence matched < 77% of a reference sequence, no taxonomic identity was assigned. To designate a sequence in a phylum the sequence was at least 77% identical to the reference sequence, for the class >80%, for the order > 85%, > 90% for the family, > 95% for the genus, and for the species >97%. We acknowledge that use of the V4 region of the 16s does not give more than a tentative evaluation of the lower phylogenetic levels (e.g. genus and species), yet we feel that some discussion of the findings in terms of genus and species provided interesting insight as part of this unique survey. Normalized to 20,000 sequences per sample, Alpha and Beta diversity analyses were completed using the Qiime2 [18] microbiome analysis pipeline (https://qiime2.org/).

## Results

We captured and sampled 10 turkey vultures, 9 bald eagles and 2 common ravens. Except for one turkey vulture captured on an Oregon coastal beach, all individuals were captured on southwest Washington coastal beaches or adjacent to Grays Harbor (Fig 1). Eight of the turkey vultures were adults and two were immatures. Three of the bald eagles were adults and six were immature; six were male and four were female. One common raven was an adult female and the other was an immature male. We collected one fecal sample from each individual for a total of 21 samples.

**Fig 1.**
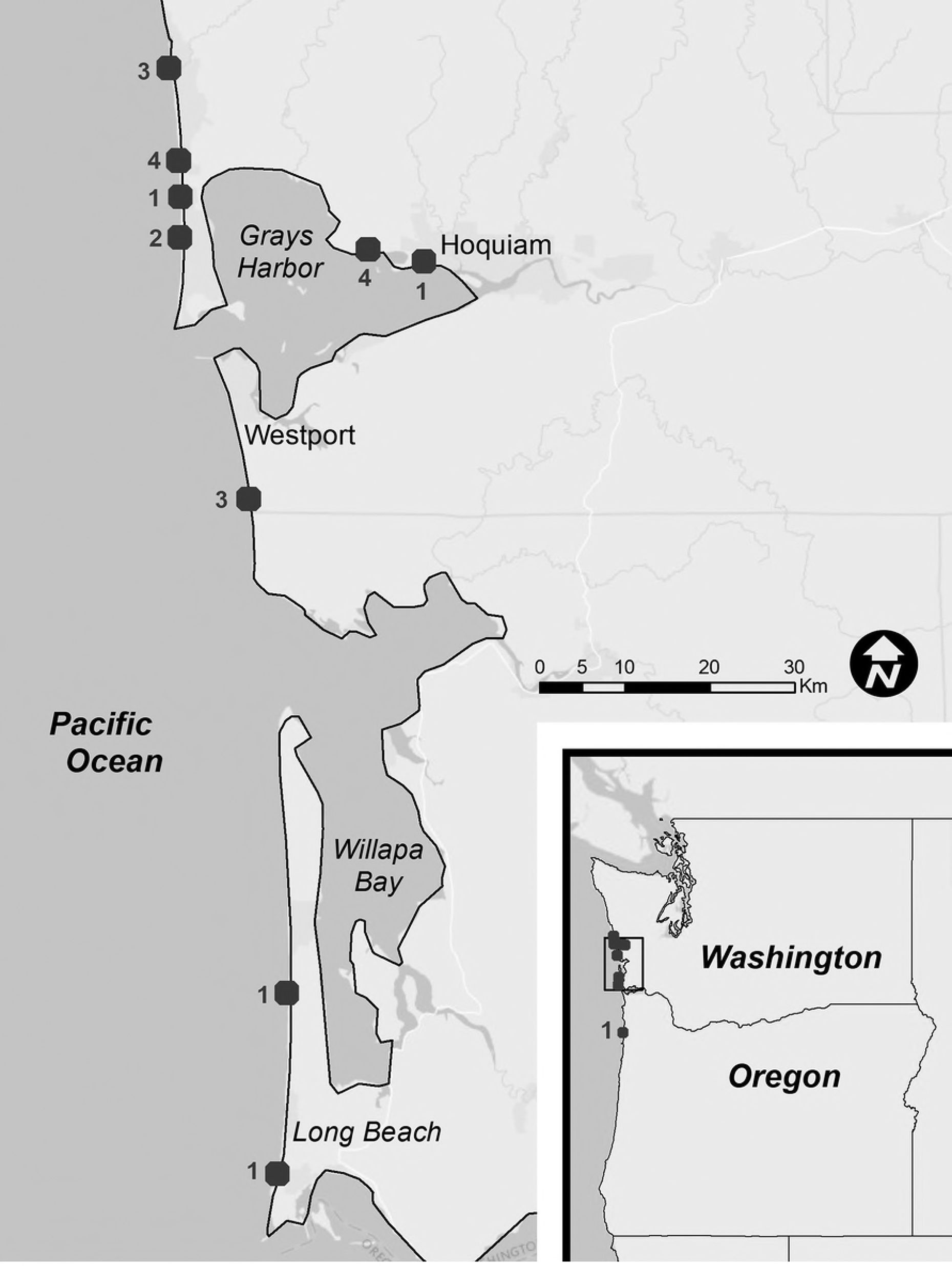
Avian scavenger sampling sites. Locations where fecal samples were collected from turkey vultures, bald eagles and common ravens in coastal Washington and Oregon. Numbers next to squares show the number of avian scavengers trapped at that location.

Genetic sequencing resulted in 1,905,510 sequences from the 21 samples we collected. Bacterial OTUs comprised 99.7% of the sequences found; 0.006% were Archaea. The other 0.25% constituted sparse cross-reactions of the primers with eukaryotes [19].

The total number of bacterial taxonomic OTUs detected among the 21 samples was 1065. Turkey vultures presented the largest diversity of bacterial fecal microbiome (1022 OTUs; *n* = 10), followed by common ravens (454 OTUs; *n* = 2), and bald eagles (604 OTUs; *n* = 9). In turkey vultures, 183 bacteria genera were identified, 141 in bald eagles, and 131 in common ravens (Table 1). Rare faction curves of observed OTUs and Shannon diversity indices were generated based on a rarefaction level of 20,000 sequences per sample (Fig 2). Non-parametric pairwise comparisons made between turkey vultures and bald eagles (Fig 3) revealed significant differences in the number of observed OTUs (p < 0.001) as well as Shannon diversity indices between the two species (p < 0.001). The alpha diversity of turkey vultures was found to be significantly greater than the alpha diversity of bald eagles with regards to both measured alpha diversity metrics. Corresponding with alpha diversity analyses, based on ANOSIM pairwise comparisons (S1 Table), beta diversity between the turkey vultures and both the bald eagles as well as the common raven is shown. Beta diversity comparisons were based on the weighted unifrac distance matrix values which can be better visualized via principal coordinate analysis (PCoA) plot (Fig 4).

**Table 1.** Number of bacteria taxonomic groups.

| Avian Species | Bacteria Taxonomy |  |  |  |  |
| --- | --- | --- | --- | --- | --- |
|  | Phylum | Class | Order | Family | Tentative Genus |
| Turkey vulture | 13 | 23 | 49 | 94 | 183 |
| Bald eagle | 12 | 22 | 43 | 78 | 141 |
| Common raven | 12 | 19 | 39 | 76 | 131 |

Number of bacteria taxonomic groups identified in fecal samples collected from turkey vultures (n = 10), bald eagles (n = 9) and common ravens (n = 2) captured on coastal beaches in Washington and Oregon.

**Fig 2.**
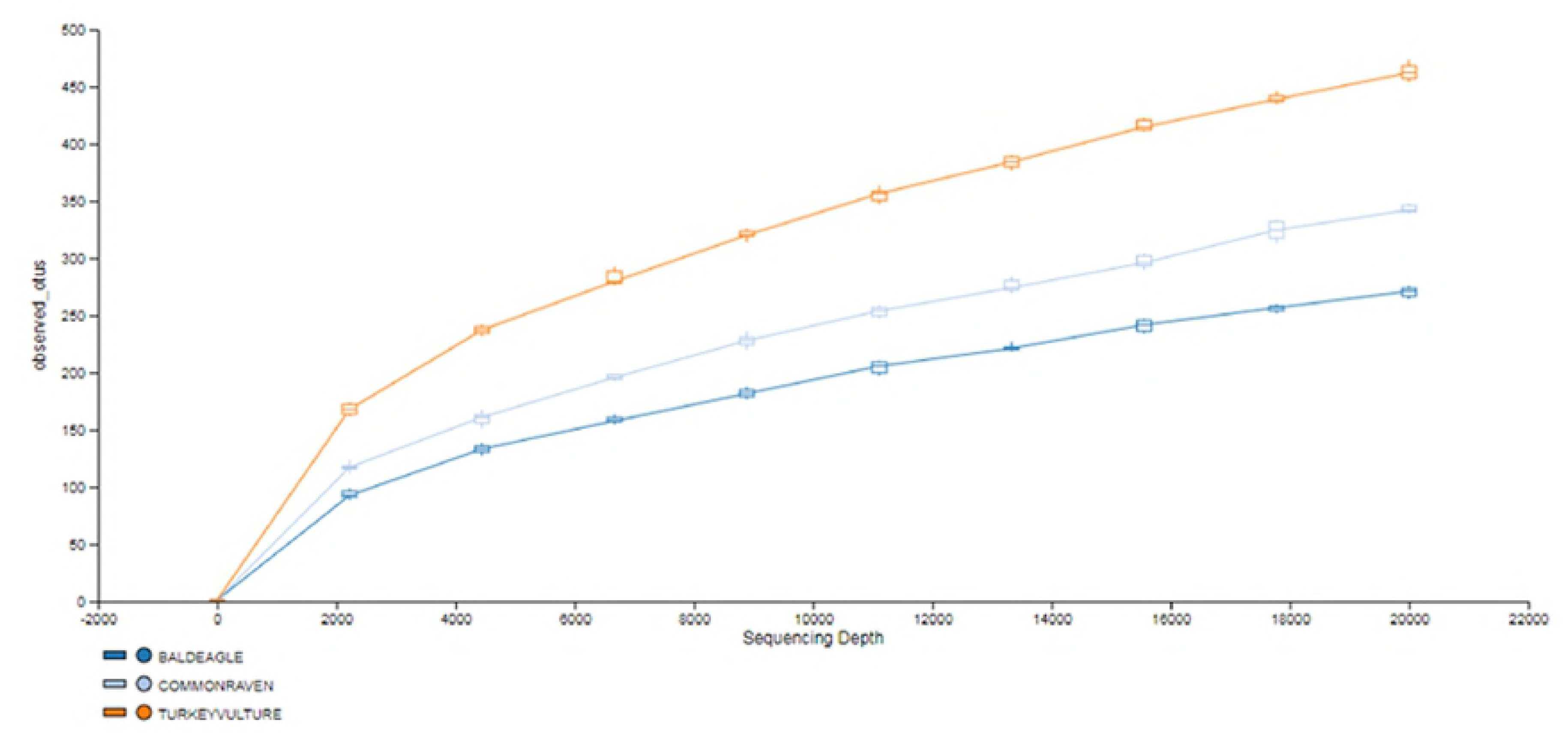
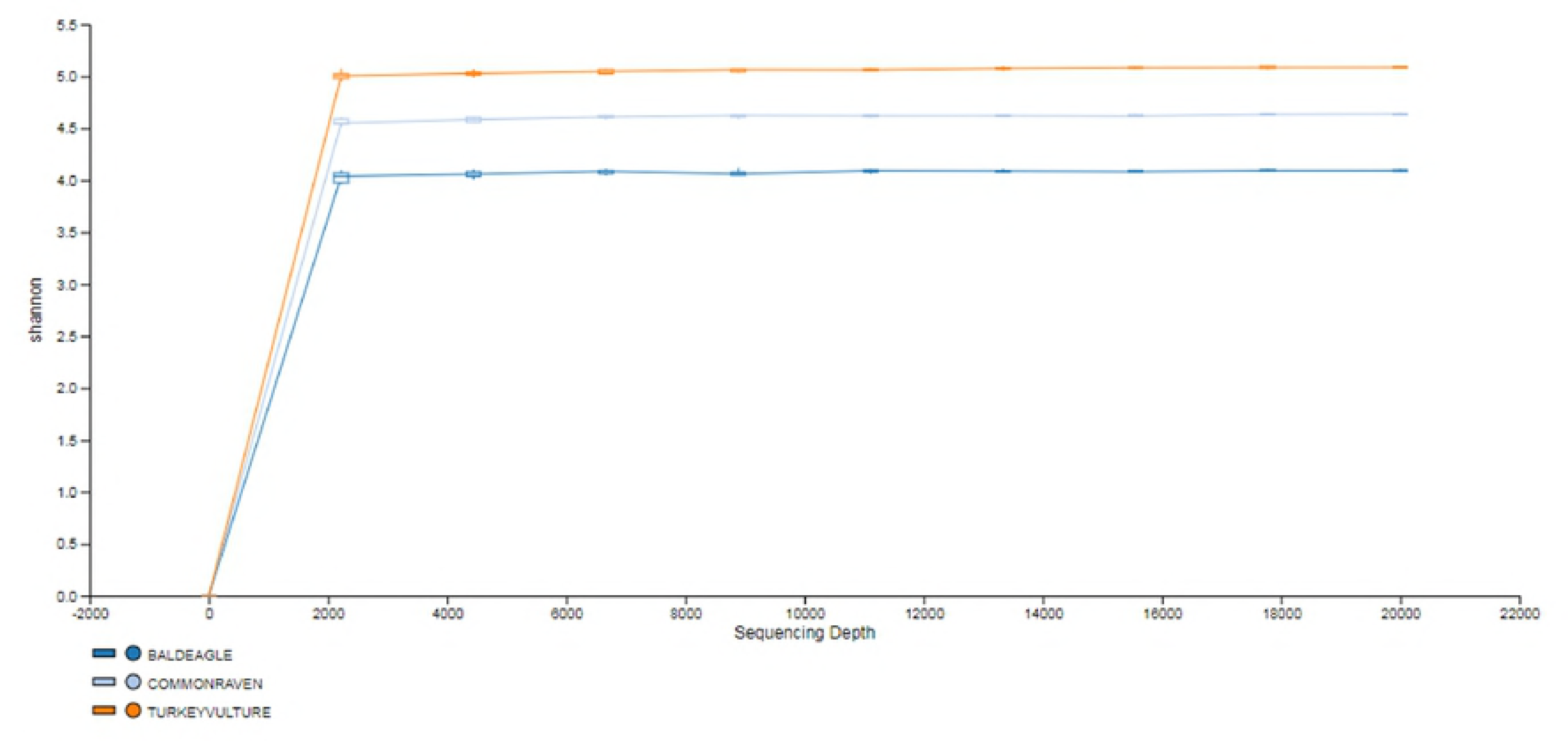
OTU Rarefaction and Shannon Diversity Index Curve. 16s rRNA sequence data was rarefied to 20,000 sequences per sample. (A) OTUs and (B) Shannon diversity indices were plotted for three avian species; Bald Eagle (n=9), Common Raven (n=2), and Turkey Vulture (n=10). OTUs were clustered at 97% identity.

**Fig 3.**
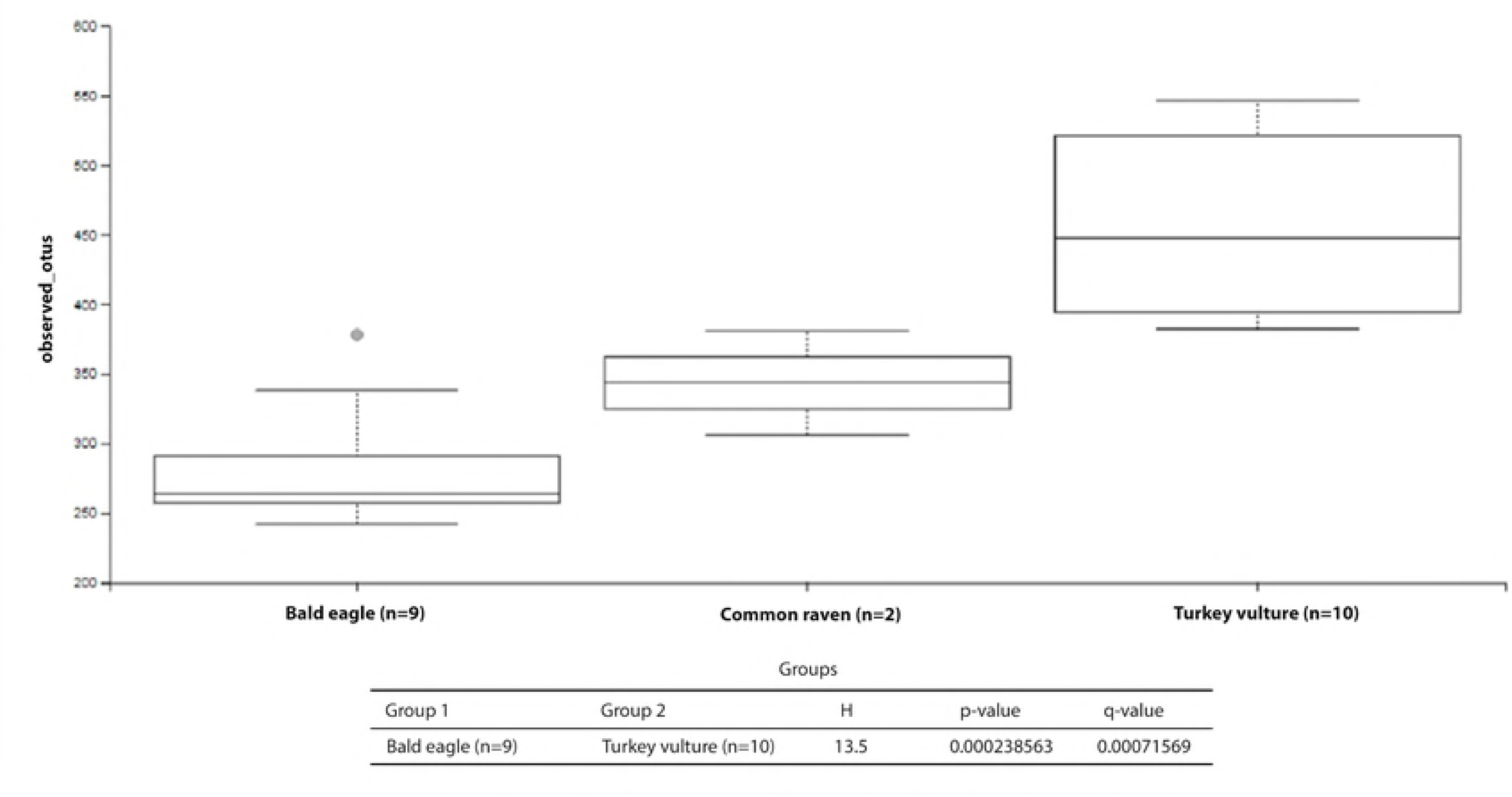
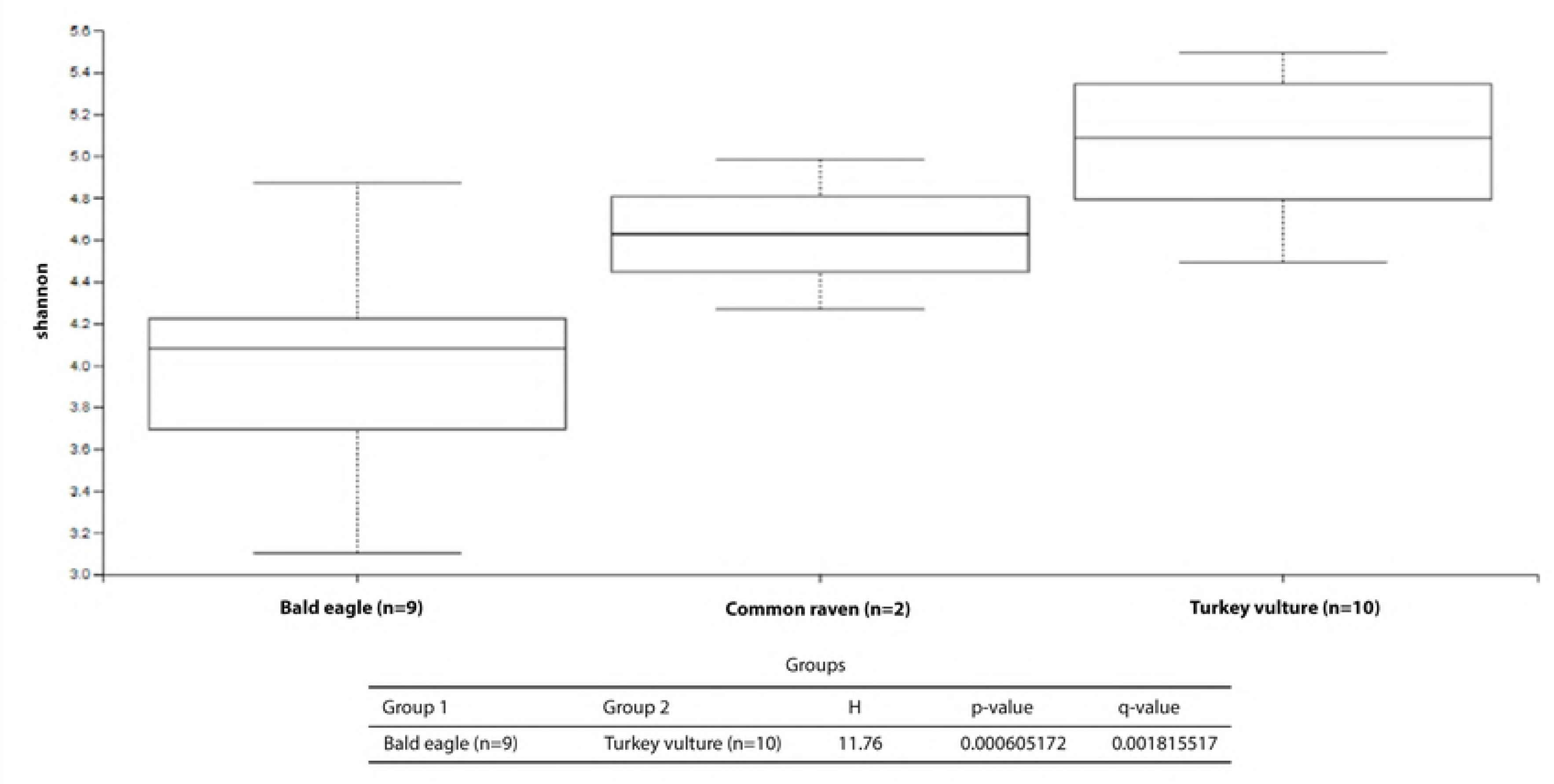
Kruskal-Wallis Pairwise Results (Observed OTUs). Significant differences present in alpha diversity metrics: (A) Observed OTUs and (B) Shannon diversity indices; were identified using the non-parametric Kruskal-Wallis H Test and differences between groups identified using multiple pairwise comparisons. Comparisons involving the Common Raven were not reported due to limited sample size (n=2).

**Fig 4.**
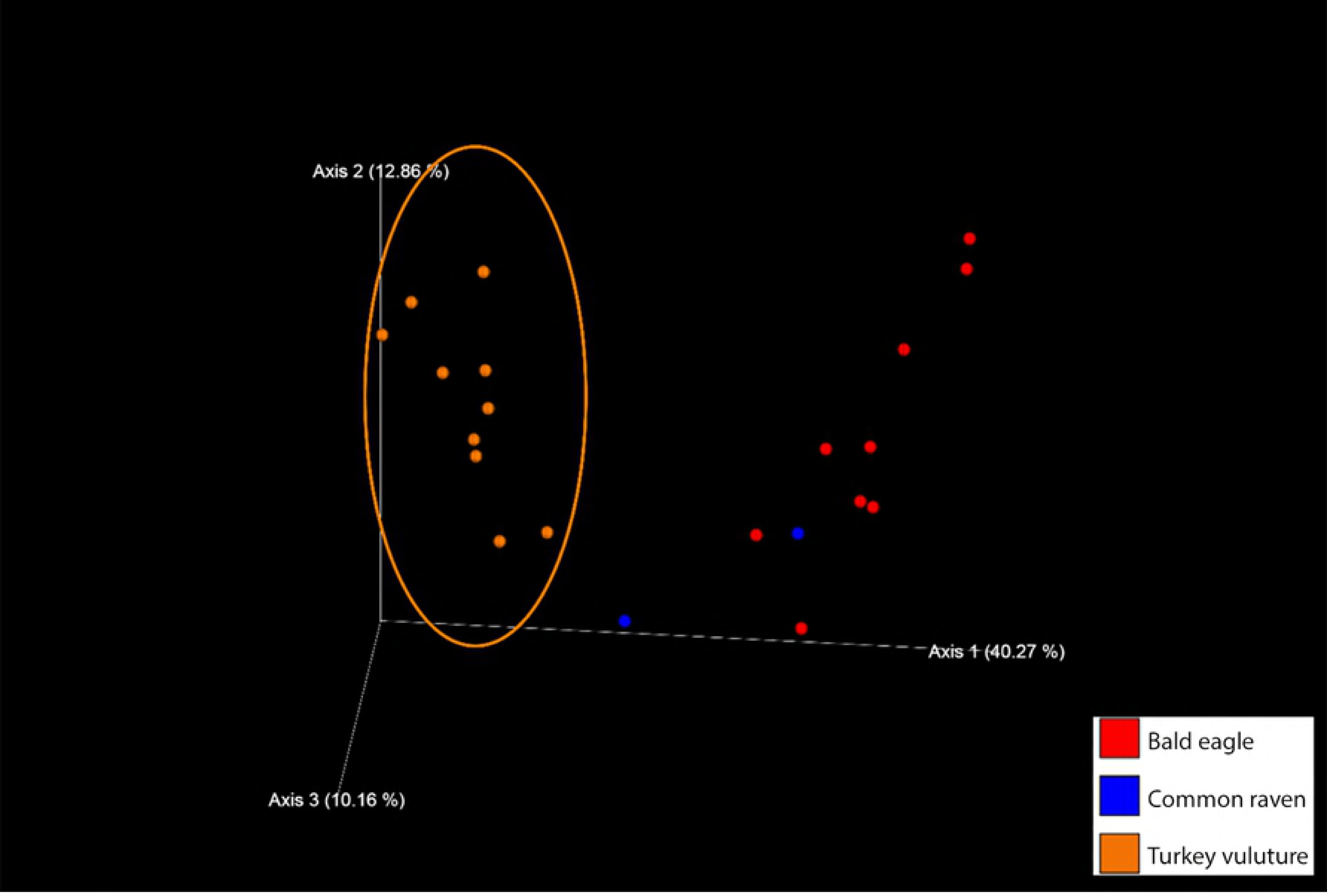
Weighted UniFrac PCoA. Principal coordinate plot of weighted UniFrac [18] data with colors keyed on the three avian species sampled. Bald Eagle (n=9; Red), Common Raven (n=2; blue), and Turkey Vulture (n=10; orange). The phylogenetic assemblage of the Turkey Vultures is clearly distinct from the remaining two sampled species.

The proportions of bacterial OTUs varied by phylum for each bird in the study (Fig 5). Thirteen phyla were represented among bacteria detected in the fecal samples (Fig 5, S1 Figure, and S2 Table). Overall, more than 80% of the bacterial sequences detected belonged to two phyla, Firmicutes and Proteobacteria. In turkey vultures, the two most common phyla were Firmicutes and Fusobacteria and in bald eagles and common ravens Firmicutes and Proteobacteria were most common.

**Fig 5.**
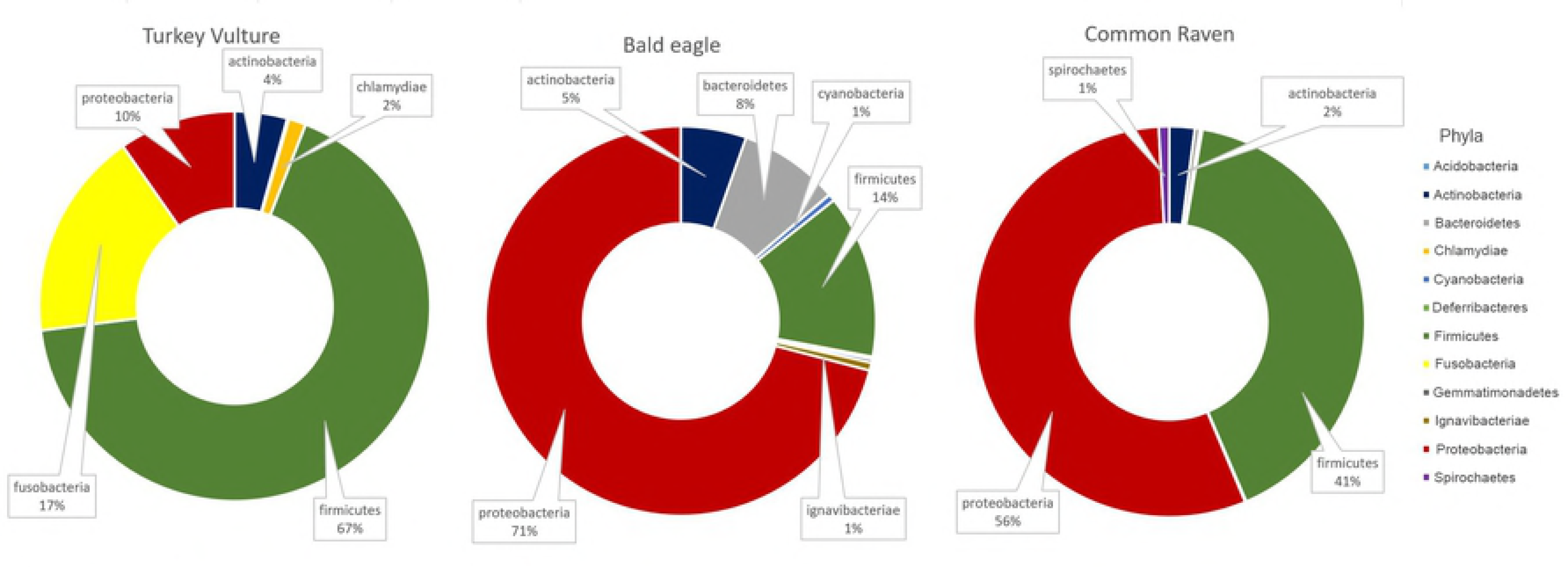
Mean proportions of bacterial operational taxonomic units by phylum. Mean proportions of bacterial operational taxonomic units by phylum detected in fecal samples collected from turkey vultures (n = 10) bald eagles (n = 9), and common ravens (n = 2) captured on coastal beaches in Washington and Oregon. Proportions < 1% are not shown.

Twenty-three classes of bacteria were represented in the fecal samples. In turkey vultures, class Clostridia contained the highest proportions of OTUs by individual, while in bald eagles class Betaprotobacteria held this distinction (Fig 6). The two common raven fecal samples also included OTUs in class Betaproteobacteria, with a larger proportion of OTUs in the sample from the adult.

**Fig 6.**
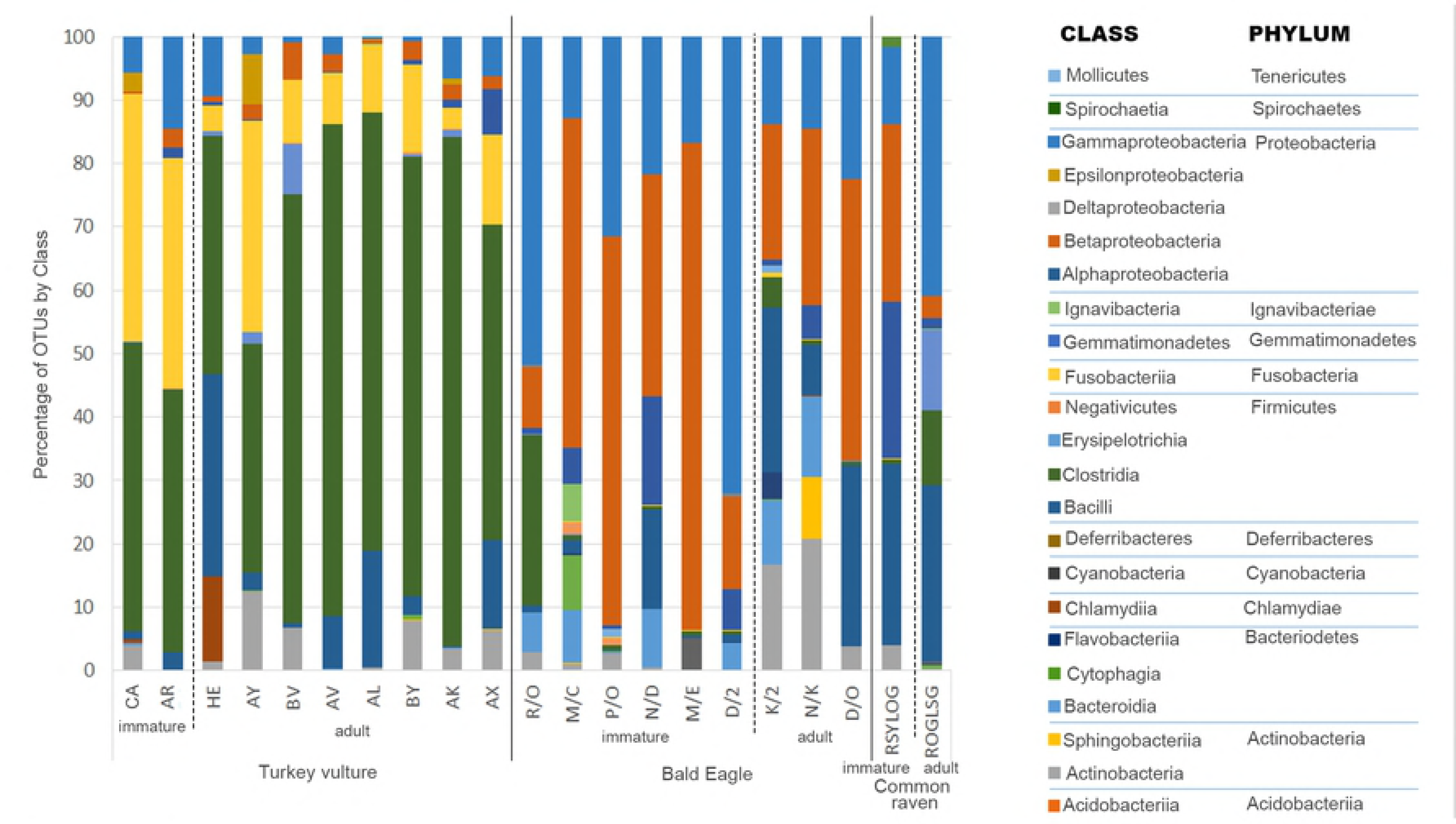
Proportions of bacterial operational taxonomic units by class. Proportions of bacterial operational taxonomic units by class detected in fecal samples from turkey vultures (n=10), bald eagles (n = 9) and common ravens (n = 2) captured on coastal beaches in Washington and Oregon. Letters below columns are the visual identification codes on bands (eagles and ravens) or wing-tags (vultures) applied to birds at capture. Proportions < 1% are not shown.

A total of 188 tentative bacterial genera were represented in the feces of the three avian species. Of the 183 different genera detected in the 10 turkey vulture samples, 18 genera accounted for 90% of the bacterial sequences, with two genera, *Clostridium* and *Fusobacterium*, comprising 48% of the total on average (Table 2). A total of 141 genera were detected in the nine bald eagle samples and 14 of these genera represented 90% of all sequences detected. The two most common, *Burkholderia* and *Pseudomonas*, comprised 36% of the total on average (Table 2). While only two common ravens were sampled, 131 bacterial genera were detected with 9 of these comprising 90% of the OTUs (Table 2). Based on hierarchal clustering of the top 35 genera detected amongst the three species, there is a clear distinction between the fecal microbiome of turkey vultures and that of the bald eagle (Fig 4).

**Table 2.**
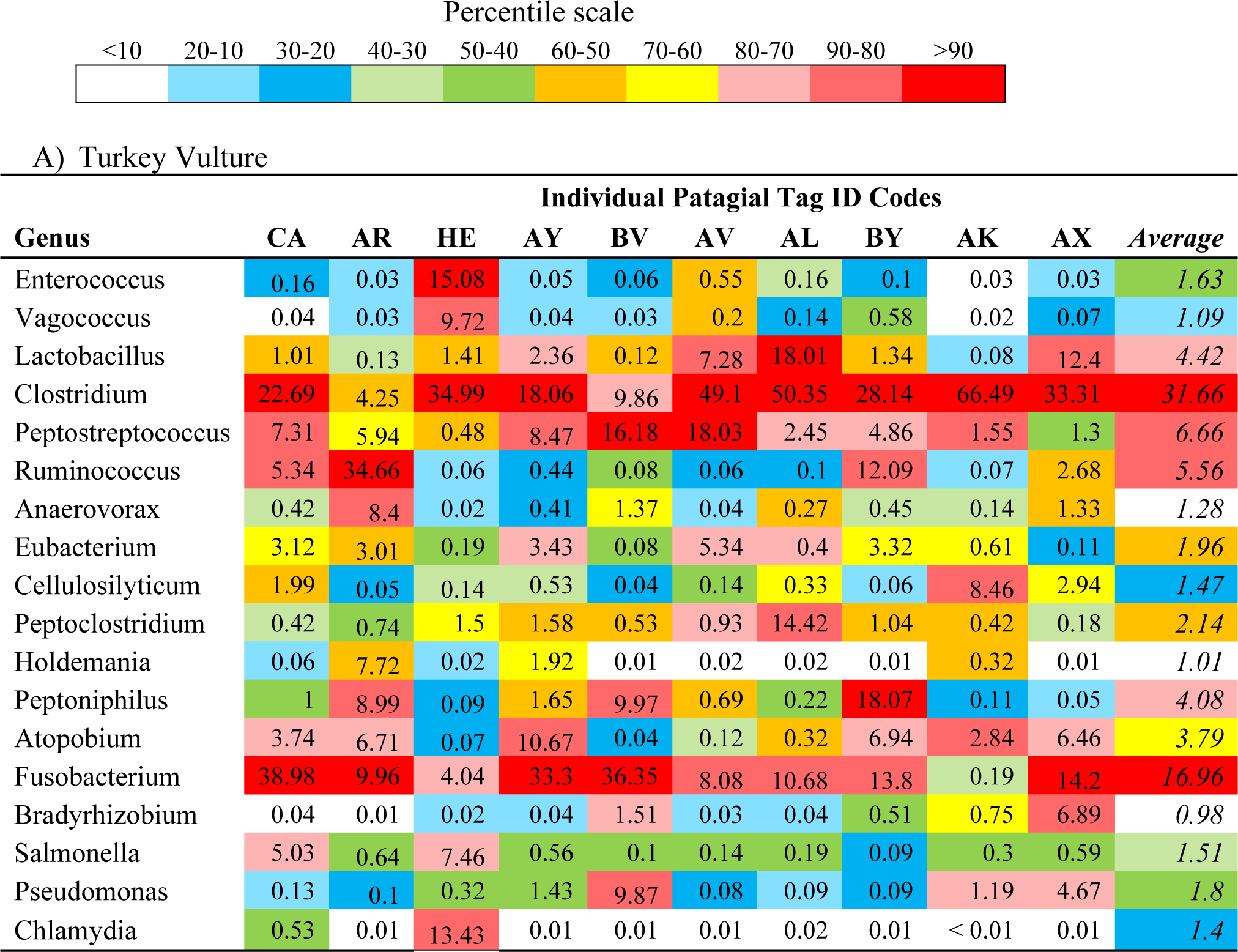
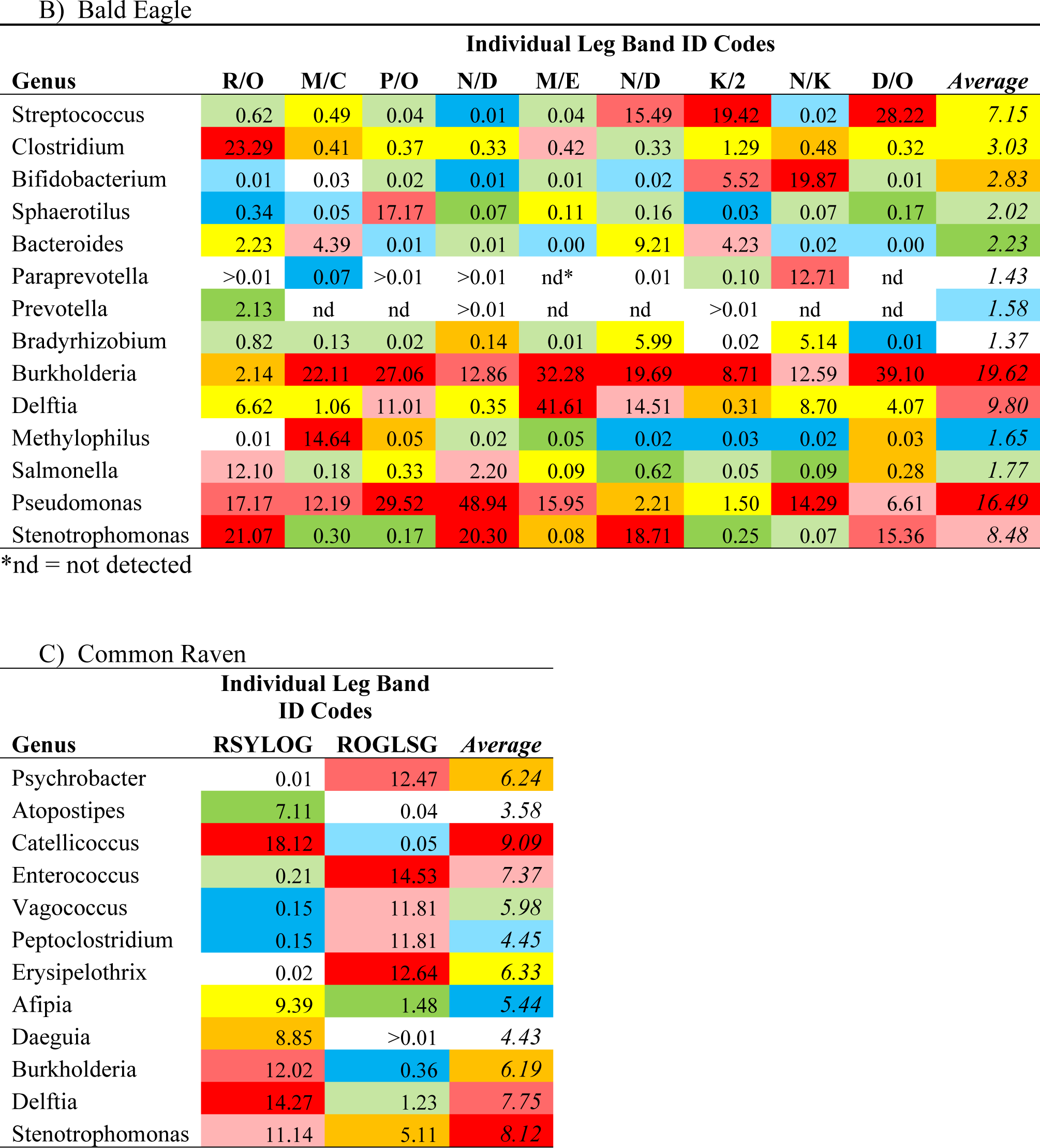
Proportions of bacterial operational taxonomic units by genus.

Top 90^th^ percentiles, expressed as a percentage of total, bacterial operational taxonomic units, by genus in fecal samples collected from A) turkey vultures (n = 10), B) bald eagles (n = 9), and C) common ravens (n = 2) captured on coastal beaches in Washington and Oregon. Letters above columns are the visual identification codes on bands (eagles and ravens) or wing-tags (vultures) applied to birds at capture. Color denotes percentile category within each column.

As with genera, most of the tentatively classified bacteria species detected across all three avian species we studied comprised relatively small proportions of the overall fecal bacterial diversity (Table 3). Moreover, bacteria that were relatively abundant in one avian species in our study were less abundant or missing altogether in the other two. In turkey vulture samples, only *Fusobacterium mortiferum*, *Clostridium perfringens*, and *Clostridium ruminantium* represented over 10% each of the total bacteria species population. In bald eagles, *Burkholderia ubonensis* comprised more than 10% of all bacterial species and *Delftia tsuruhatensis* comprised nearly 10%. In contrast, no bacteria species represented ≥ 10% of the total bacteria in common ravens.

**Table 3.**
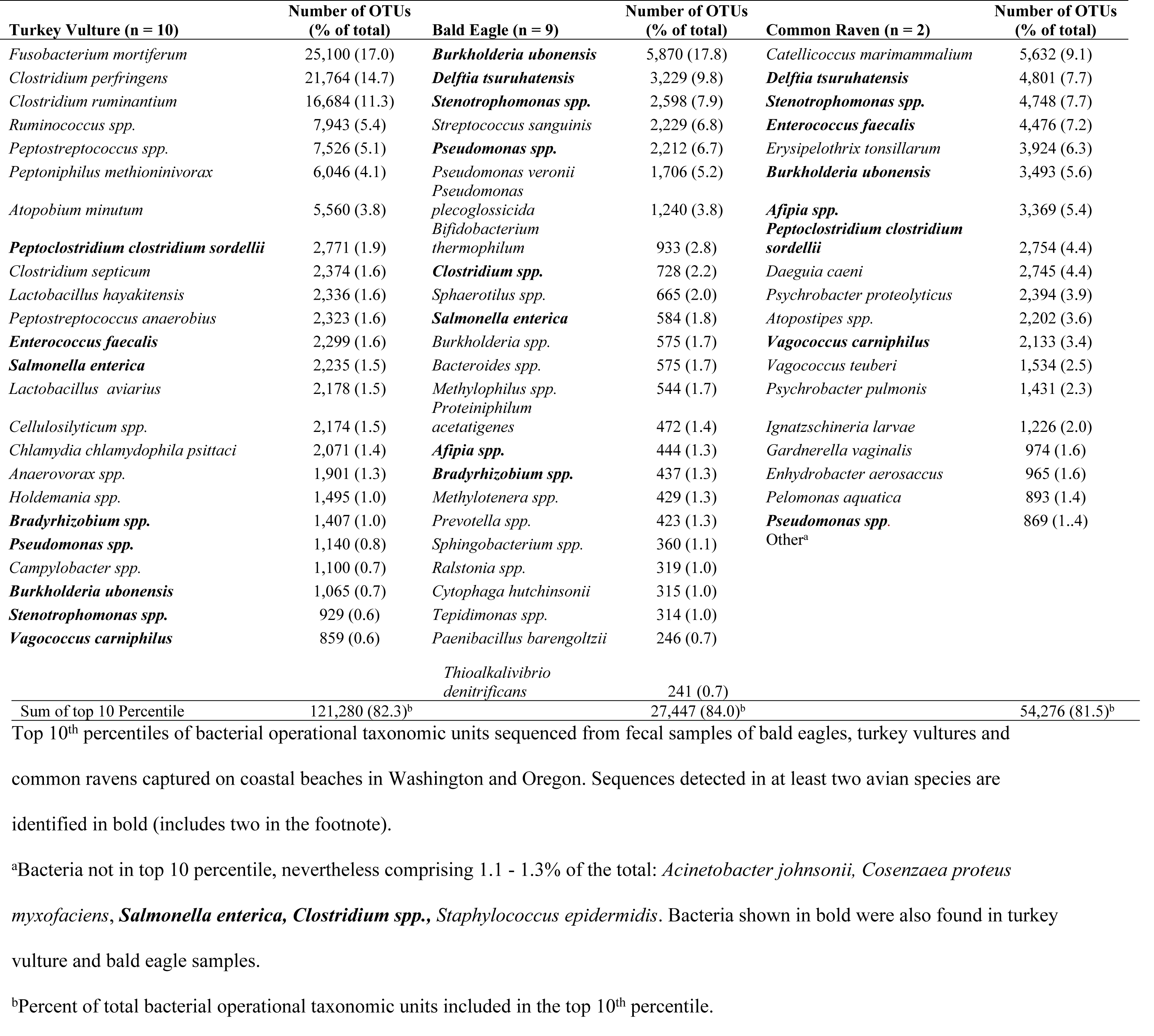
Top 10^th^ percentiles of bacterial operational taxonomic units tentatively classified to the closest species match in fecal samples of turkey vultures, bald eagles and common ravens.

Archaea were identified in one turkey vulture sample and in two bald eagle samples. We tentatively classified these as uncultured *Candidatus nitrosocaldus.* No Archaea were identified in common ravens.

## Discussion

Our analyses showed substantial microbiome diversity among turkey vultures, bald eagles, and common ravens. These differences probably represent their diverse anatomies, physiologies, and also to an extent, differences in diet. Turkey vultures, bald eagles and common ravens are among the many terrestrial vertebrates that scavenge opportunistically on the Pacific Northwest coast [20] and also inland from the coast. It should be noted that ceca, which act as sites of anaerobic activity in some species of birds, are vestigial in all three of our study species [21].

The authors are aware that bacteria species classification based upon V4 is tentative. The classification of ribosomal RNA genes has been the gold standard for molecular taxonomic research for decades, but standard primers for bacterial species identification does not exist yet [22]. On the other hand, sequences of the most commonly identified bacterial genera and species (those that are the 10 top percentile) in this study exist in the databases. These bacteria were not environmental or novel bacteria. For this reason, we discuss not only bacteria class, order, and phylum, but also genus and species.

Two classes, Fusobacteria and Clostridia, which are comprised exclusively of anaerobes, predominated the cloacal flora in turkey vultures. Roggnbuck et al. [23] also reported these two classes as most prevalent in turkey vultures. *Fusobacterium mortiferum*, which on average comprised 17% of all bacterial species detected in Turkey Vulture feces, is a normal inhabitant of the alimentary tract in chickens where it protects against pathogenic bacteria [24].

Many clostridia species are normal gut flora and play a crucial role in gut homeostasis, however under stressful gut conditions (i.e. inflammation, parasitism, etc) these bacteria may release toxins that can cause serious diseases in humans and other animals, including birds [25, 26]. *Clostridium perfringens* and *C. ruminantium* accounted for 26% of the bacterial species we detected in turkey vulture feces in our study. *C. perfringens* is known to be a normal component of intestinal flora, but can cause necrotic enteritis in birds when intestinal dysbiosis occurs [27]. *C. ruminantium* is reported to contribute to rumen fermentation in cattle and it is also a food-spoiling bacterium, but it has not been directly linked to animal diseases [28].

Bacterial toxins can also be present in carrion. Turkey vultures have a remarkable tolerance for some of these toxins, such as botulinum [29, 30] and anthrax [23, 31]. Only 0.3% of sequences assembled from turkey vultures feces were identified as *C. botulinum* and no *Bacillus anthracis* was detected in these birds. Neither of the aforementioned species was present in bald eagles or common raven samples.

Most *Enterococcus* (phylum Firmicutes) are commensal bacteria; however there are a few species in this genus, including *E. faecalis*, that are associated with significant mortality in humans and other animals [32, 33]. *Enterococcus faecalis* comprised 7.2 % of the bacterial OTUs in the two common ravens we sampled and 1.6% in the turkey vultures. While this bacterium was detected in all nine bald eagle fecal samples, they comprised < 0.1% of the assembled sequences for that species. In Illinois, 25 raptors were tested for *Enterococcus* due to their potential to develop resistance to antibiotics; 53 of 56 cloacal samples collected contained *E. faecalis* [34]. *Streptococcus sanguinis* (phylum Firmicutes) comprised nearly 7% of all bacteria in bald eagles but ≤ 0.3% in turkey vultures and common ravens, Although these bacteria are considered normal inhabitants of mammals and humans [35], there is no information about the commonality of these bacteria or their significance in the digestive tracts of avian species. *Erysipelothrix tonsillarum* comprised 6% of the OTUs in the two common raven fecal samples and < 0.1% in turkey vultures and bald eagles. These non-pathogenic bacteria are commonly isolated from healthy swine and poultry as well as poultry litter. *E. tonsillarum* is closely related to the pathogenic species *E. rhusiopatheae*, causative agent of erysipeloid in mammals and erysipelas in poultry [36, 37].

Almost 70% of fecal bacteria in the bald eagles we sampled (n = 9) and 55% in the two common ravens belonged to the phylum Proteobacteria. In contrast, Proteobacteria accounted for less than 10% of the turkey vulture microbiome (n = 10). In bald eagles, the most common species belonging to this phylum were tentatively classified as *Burkholderia ubonensis* and *Delftia tsuruhatensis*, comprising 18% and 10% of all fecal bacteria detected respectively. Both are common environmental bacteria that have been occasionally linked to infections in humans [38, 39]. *B. ubonensis* was also detected in the two common ravens sampled, ranking seventh most common and averaging 5.6%. *Catellicoccus marimammalium* was the most common bacteria in the common ravens sampled, averaging 9% of the total. *C. marimammalium* was the most common species in the feces of gulls in Wisconsin, U.S.A. [40].

Other notable bacteria detected from the Proteobacteria phylum included bacteria in *Escherichia*, *Campylobacter*, *Pseudomonas*, and *Salmonella*. *E. coli* inhabits the large intestine and distal ileum of most vertebrate species, including birds; the organism is shed in the feces. Generally-speaking, the vast majority of the strains are components of normal intestinal flora and are non-pathogenic. Some strains are of low virulence and may cause opportunistic infections. A few are highly pathogenic and may cause septicemia. *E. coli* sequences were detected in the cloacal samples from eight of the ten turkey vultures, eight of the nine bald eagles, and both ravens. However, *E. coli* sequences were sparse and, on average, accounted for less than 0.01% of all the bacterial OTUs per bird. No further analysis was done on these *E. coli* sequences to determine if they had any pathogenic genes or not. *Campylobacter* sp. were detected in all the fecal samples from turkey vultures and accounted for 0.7% of the bacterial OTUs (Table 3). In bald eagles and common ravens, *Campylobacter* sp. were also detected in all samples, but the OTU frequency was less than 0.01%. Members of *Pseudomonas* are ubiquitous and can be opportunistic pathogens [41]. Bacteria from the *Pseudomonas* genus were detected in all fecal samples. They constituted 17% of the fecal flora in bald eagles, but < 2% of all bacterial OTUs in turkey vultures and common ravens.

Avian predators may become infected with *Salmonella* when they consume infected animals [42]. *S. enterica* colonize the intestinal tract and have been isolated from wild birds with or without clinical signs of disease [43]. In our study, *S. enterica* comprised 1-2% of bacterial species detected in all three avian scavengers In other research, *S. enterica* sampled from the feces of scavenging birds ranged from 1% to 20% by isolation [44–46]. Differences in prevalence between isolation and detection by meta-analysis may be due to the stability of the DNA molecule that can be amplified even when the target organism is no longer viable. *S. enterica* sequence was performed only to the species level, and subspecies or serotype were not determined.

Archaea are single-celled prokaryotes that are difficult to culture [47]. They have been found in the digestive tracts of some animals where they produce methane [48]; the role these organisms play in the gut of birds is still unknown [1]. *Candidatus nitrosocaldus* is ubiquitous in water environments being present in plankton, sediments, and surrounding soils of lakes and oceans [49]. It has a particular affinity for hot spring environments. It is also common in waste water treatment plants and fertilized soils. Its presence in the feces of two bald eagles and one turkey vulture in our study may be a reflection of their contact with marine carrion.

In summary, this is the first study utilizing metagenomics to investigate the fecal microbiome to the species level. This is also the first meta-analysis performed in bald eagles and common ravens. We demonstrated that the fecal flora varies among the three scavenger species even though they live in the same region and consume some of the same foods (D. Varland, unpub. data). We discovered that over 50% of the microbiota in the cloaca belongs to a single phylum in all three species, Proteobacteria for bald eagles and common ravens and Firmicutes for turkey vultures. As expected numerous species of microorganisms were tentatively identified; however, only about 10% of these tentative species contained > 1% of the OTUs. Finally, this method for identification and characterization of microorganism populations can be used in epidemiological, pathogen detection, and microbial diversity studies.

## Acknowledgements

We thank the Washington State Parks and Recreation Commission and the Oregon Parks and Recreation Department for allowing vehicle access to coastal beaches, which was essential to our work. We also greatly appreciate the field assistance of many, including Nathalie Denis, Glenn Marquardt, Larry Warwick, Sandra Miller, Pam McCauley, Suzy Whittey, Tom Rowley and Dale Larson. D. Varland color marked birds in the study under federal banding permit 21417 and under state Scientific Collection Permits13-014, 14-084 and 15-093 in Washington and under Scientific Taking Permit 049-13 in Oregon.

## Supporting information

**S1 Figure. Proportions of bacterial operational taxonomic units by phylum.** Proportions of bacterial operational taxonomic units by phylum detected in fecal samples from turkey vultures (n = 10), bald eagles (n = 9), and common ravens (n = 2) captured in coastal Washington and Oregon. Letters below columns are the visual identification codes on bands (eagles and ravens) or wing-tags (vultures) applied to birds at capture. Proportions < 1% are not shown.

**S2 Figure. Hierarchal Clustering of Taxonomic Data.** Evaluation of the taxonomic classification data using a dual hierarchal dendrogram. Each sample is clustered on the X-axis labeled based on the sampled species. Samples with more similar microbial populations are mathematically clustered closer together. The heatmap represents the relative percentages of each genus. The predominant genera are represented along the right Y-axis. The legend for the heatmap is provided in the upper left corner. The microbial population of Turkey Vultures (n=10) is distinct from the population identified in the Bald Eagle (n=9).

**S1 Table.** Pairwise ANOSIM Results (Weighted UniFrac) Significant differences present in the weighted UniFrac distance matrix values were identified using the Analysis of Similarities (ANOSIM) Test and differences between groups identified
 using multiple pairwise comparisons over 999 permutations.

| Group 1 | Group 2 | Sample size | R<sup>a</sup> | p-value | q-value |
| --- | --- | --- | --- | --- | --- |
| BALDEAGLE | COMMONRAVEN | 11 | 0.204204 | 0.239 | 0.239 |
| BALDEAGLE | TURKEYVULTURE | 19 | 0.928121 | 0.001 | 0.003 |
| COMMONRAVEN | TURKEYVULTURE | 12 | 0.81087 | 0.015 | 0.0225 |

*<sup>a</sup>R scales from +1 to -1. A value of +1 indicating the most similar samples are in the same* *group. A value equal to 0 indicating there is no relationship observed between similar and* *dissimilar samples.*

**S2 Table.** Mean number (and percent) of bacterial operational taxonomic units by phylum. Mean number (and percent) of bacterial operational taxonomic units by phylum detected in fecal samples from turkey vultures (n = 10), bald eagles (n = 9), and common ravens (n = 2) captured on coastal beaches in Washington and Oregon. For each species, phyla with the largest number of operational taxonomic units are identified in bold.

| Phylum | Turkey Vulture |  | Bald Eagle |  | Common Raven |  |
| --- | --- | --- | --- | --- | --- | --- |
|  | Mean | (Percent) | Mean | (Percent) | Mean | (Percent) |
| Acidobacteria | 5.7 | (< 0.1) | 0.2 | (< 0.1) | ND<sup>a</sup> | ND |
| Actinobacteria | 5441 | (4.3) | 2,033.8 | (5.4) | 951 | (2.1) |
| Bacteroidetes | 261.4 | (0.2) | 2,912.2 | (8.3) | 336.5 | (0.4) |
| Chlamydiae | 2,000 | (1.4) | 6.9 | (< 0.1) | 5.5 | (< 0.1) |
| Cyanobacteria | 7.3 | (< 0.1) | 195.3 | (0.6) | 150.5 | (0.2) |
| Deferribacteres | 19.2 | (< 0.1) | 0.1 | (< 0.1) | ND | ND |
| Firmicutes | **10,6044.8** | **(67.1)** | **4,937.4** | **(13.6)** | **27,467** | **(40.9)** |
| Fusobacteria | **22,873.3** | **(17.3)** | 81.2 | (0.2) | 61.5 | (0.1) |
| Gemmatimonadetes | 4.1 | (< 0.1) | 111.4 | (0.3) | 1 | (< 0.1) |
| Ignavibacteriae | 7.9 | (< 0.1) | 208.3 | (0.6) | 1 | (< 0.1) |
| Proteobacteria | 11,345.5 | (9.7) | **22,445.3** | **(70.9)** | **32,631** | **(55.5)** |
| Spirochaetes | 2.4 | (< 0.1) | 0.3 | (< 0.1) | 372.5 | (0.9) |
| Tenericutes | 2.1 | (< 0.1) | ND | ND | ND | ND |

<sup>a</sup> ND = Not detected.

## Abbreviations

OTU: operational taxonomic unit

